# Foxm1 drives cardiomyocyte proliferation in adult zebrafish after cardiac injury

**DOI:** 10.1101/2022.06.22.497073

**Authors:** Daniel A. Zuppo, Maria A. Missinato, Lucas Santana-Santos, Guang Li, Panayiotis V. Benos, Michael Tsang

## Abstract

The regenerative capacity of the mammalian heart is poor with one potential reason being that adult cardiomyocytes cannot proliferate at sufficient levels to replace lost tissue. During development and early neonatal stages, cardiomyocytes can successfully divide under injury conditions; however, as these cells mature their ability to proliferate rapidly decreases. Therefore, understanding which regulatory programs are required to induce post-mitotic, mature cardiomyocytes into a proliferative state is essential in order to enhance cardiac regeneration. Unlike mammals, adult zebrafish cardiomyocytes in the injury border zone do proliferate. This model provides an opportunity to elucidate how these border zone cells respond to different stimuli post-injury and to study which regulatory programs are required for adult cardiomyocyte proliferation. Here we report the forkhead transcription factor, *foxm1*, is required for cardiomyocyte proliferation after cardiac injury through transcriptional regulation of cell cycle genes. Transcriptomic analysis of injured adult zebrafish hearts revealed that *foxm1* expression is increased after injury in border zone cardiomyocytes. *foxm1* mutants showed decreased cardiomyocyte proliferation after ventricular resection, resulting in larger fibrotic scars. Moreover, decreased expression of cell cycle progression genes suggests that Foxm1 is required for different cell cycle checkpoints during cardiomyocyte division. Subsequent analyses of Foxm1 targets revealed the microtubule and kinetochore binding protein, *cenpf*, is required for cardiac regeneration as *cenpf* mutants failed to regenerate due to increased cardiomyocyte binucleation. Thus, *foxm1* and *cenpf* are required for cardiomyocytes to complete mitosis during zebrafish cardiac regeneration.

## Introduction

Cardiac regeneration is a phenomenon where multiple cell types respond after injury to replace damaged heart tissue. Urodele amphibians [1, 2], zebrafish [3-6], and neonatal mice [7] possess this regenerative capacity, whereas others, such as medaka [8] and adult mammals [9], lack this ability. A major difference between regenerative and non-regenerative hearts is whether cardiomyocytes divide after injury to replenish the damaged tissue. While some species retain this ability throughout their lifespan [1-6], others possess a limited proliferative window that is lost as cardiomyocytes mature. In mice, neonatal cardiomyocytes can robustly divide after injury, but division rapidly decreases after post-natal day 7 (P7) and fibrotic tissue persists at the injury site [7]. Understanding how this proliferative window is determined would allow us to stimulate cardiomyocyte proliferation in non-regenerative, injury settings.

Embryonic cardiomyocytes activate multiple pathways that promote maturation within these cells. These processes include changes in sarcomere assembly, metabolism, and cell cycle activity. Loosely organized myofibrils are produced in embryonic cardiomyocytes at the beginning of sarcomere assembly, but a large expansion in myofibril number occurs during cell maturation coupled with increased organization of the sarcomere into banded, contractile structures of the Z-line and M-line [10-12]. In addition to this, mouse embryonic cardiomyocytes produce sarcomere components such as *Myh7, Myl7*, and *Tnni1* while adult cells predominantly express *Myh6, Myl2*, and *Tnni3* after maturation [11-17]. With increased contractile organization, adult cardiomyocytes also require higher levels of ATP to sustain cardiac function. Mature cardiomyocytes use fatty acid oxidation to generate substrates required ATP generation via oxidative phosphorylation [18]. These cells have increased mitochondria number, size, and more developed cristae which support the high levels of ATP required for contraction in adult cardiomyocytes [19]. In contrast, embryonic cardiomyocytes are reliant on glycolysis as their primary energy source which is shown through increased expression of the glycolysis initiation gene Hexokinase 1 (*Hk1*) [20] and the presence of immature mitochondria [21]. Cell cycle activity is another hallmark that indicates the maturation status of cardiomyocytes as they switch from a proliferative state to embryonic cells and exit the cell cycle as they mature. During mouse development, cardiomyocyte proliferation peaks at embryonic day 12 (E12) and P4-6. These cells have increased mRNA and protein levels of cyclins and cyclin-dependent kinases (CDKs) required for cell cycle progression [22-24]. However, the expression of cyclin-dependent kinase inhibitors (CKIs) increases by P3-P6 and remains elevated in cardiomyocytes, preventing cell cycle progression [22, 23, 25, 26]. Concurrent with the reduction in cycling cardiomyocytes, the ploidy type begins to switch from mononucleated diploid to either a mononucleated or binucleated polyploid cells [23, 24, 27, 28]. Taken together, this suggests that multiple regulatory programs coordinate to initiate cardiomyocyte maturation and attenuate the proliferation of these cells. Once cardiomyocytes mature in mammals, it becomes increasingly difficult to drive them into a regenerative state and this is one of the reasons why fibrotic tissue remains after cardiac injury.

Unlike mammals, adult zebrafish cardiomyocytes can dedifferentiate from a mature to an embryonic state and exhibit robust proliferation under different injury conditions [4-6, 12, 29]. This makes them an ideal model to study which genes regulate the proliferative switch in these cells. In the cardiac resection model, 20% of the ventricle is surgically removed and a clot forms at the site of injury [5]. Cardiomyocytes respond to induced stimuli at 3 days post-amputation (dpa) and begin to dedifferentiate and divide. By 7dpa, cardiomyocyte proliferation peaks and decreases by 14dpa. Using this model, studies have shown that cardiomyocyte proliferation can be activated via multiple factors [30-33]. However, many of these genes encode secreted ligands involved in cell non-autonomous activation of the cell cycle and the identification of downstream transcription factors required for cardiomyocyte proliferation is not well characterized.

In this study, RNA-sequencing was performed on uninjured and resected hearts to identify genes involved in mitotic regulation. Forkhead box M1 (*foxm1*), a transcription factor, was significantly increased at 3dpa and was deemed a candidate for injury-induced cardiomyocyte proliferation based on its known role in driving proliferation in cancer [34-40]. Indeed, we show that *foxm1* is expressed in a subset of cardiomyocytes within the injury zone and that *foxm1* mutant hearts displayed significantly decreased cardiomyocyte proliferation. Transcriptome analysis of *foxm1* mutants revealed decreased expression of AP-1, glucose metabolism, and G_2_/M phase cell cycle genes, indicating impaired cardiomyocyte proliferation. Of these G_2_/M genes, centromere protein F (*cenpf*), a canonical target of Foxm1, is required for the completion of chromatin segregation, and its depletion induces mitotic arrest in mammalian cells [41-44]. *cenpf* mutant hearts accumulate more binucleated cells, revealing a failure to complete mitosis. This study demonstrates the importance of Foxm1 as a regulator of cell cycle progression but also reveals a role for Cenpf in cardiomyocyte mitosis during heart regeneration.

## Results

### Identification of mitotic genes involved in cardiac regeneration

We previously determined that Dusp6, an ERK phosphatase, limits heart regeneration in zebrafish [45]. Heart regeneration in *dusp6* mutants was accelerated due to increased cardiomyocyte proliferation and neovascularization. To investigate transcriptome changes during the early stages of cardiac regeneration, we analyzed differential gene expression (DGE) in adult hearts collected from uninjured or injured WT and *dusp6* mutant zebrafish. Ventricular amputated hearts were extracted at 3dpa, and total RNA was isolated for RNA-sequencing (RNA-seq). For analysis, WT and *dusp6* mutant data were combined based on their injury status. DGE analysis revealed that 1149 genes were significantly increased while 153 genes were decreased at 3dpa (**Fig. 1A**, and **Supplementary Table 1**). Many genes identified in this cluster, such as *ect2, plk1*, and *hmmr* [46-48], are known to play critical roles in zebrafish cardiac regeneration and detecting these transcripts demonstrated that these injured hearts were in a proliferative state (**Supplementary Table 1**). Consistently, NIH DAVID showed the enrichment of cell cycle and DNA-binding genes using functional annotation clustering (FAC) (**Fig. 1B, and Supplementary Table 2**). To validate the RNA-seq data, qPCR was performed using uninjured and 3dpa hearts to confirm that candidate genes were increased upon injury (**Fig. 1C**). qPCR revealed increased expression of DNA-binding factors (*alx4a, foxm1, pbx3b*) and chromatin modifiers (*kmt5ab* and *hdac7b*) but not stemness factors (*pou5f3* and *zeb2b*) (**Fig. 1C**).

**Figure 1:**
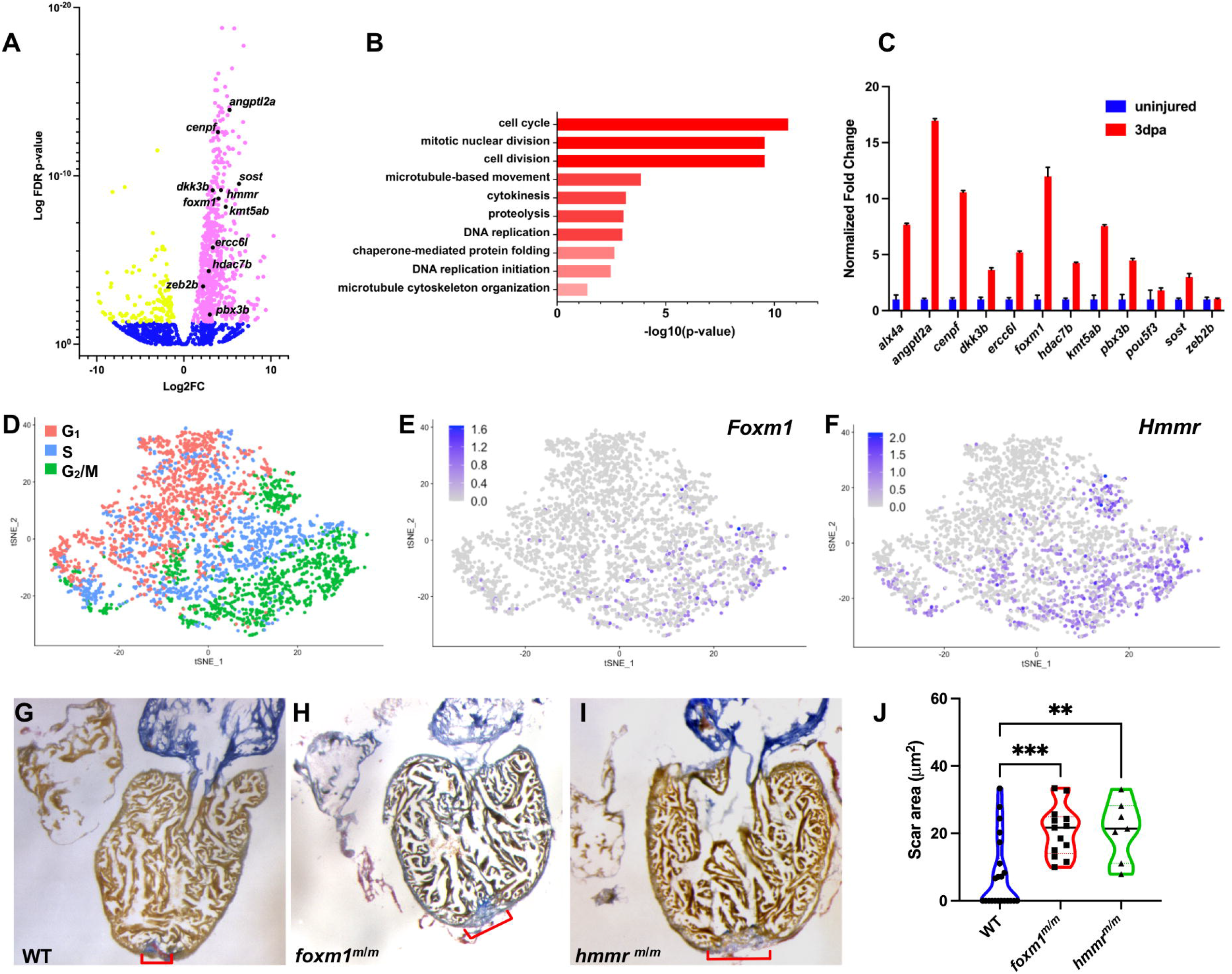
Identification of mitotic genes involved in cardiac regeneration. Representative volcano plot of RNA-seq from uninjured vs 3dpa adult ventricles (**A**). Magenta denotes genes with increased transcripts at 3dpa, yellow represents decreased transcripts, and blue represents gene with no significant change. Genes with increased transcription were analyzed with Functional Annotation Clustering (NIH DAVID) and biological processes were ranked by z-score (**B**). qPCR validation was performed on candidate genes and the representative image shown depicts fold change + S.D. (**C**). scRNA-Seq showed E10.5 embryonic left ventricular cardiomyocytes sorted by cell cycle phase (**D**). *Foxm1* (**E**) and *Hmmr* (**F**) were present in G2/M phase cardiomyocytes. Representative images from WT 30dpa (**G; n=22**), *foxm1*^*m/m*^ 30dpa (**H; n=13**), and *hmmr*^*m/m*^ 30dpa (**I; n=7**) hearts stained with AFOG. AFOG staining labels fibrin (red), collagen (blue), and muscle (orange-brown). Scar area was calculated as pixel value and converted into microns squared (μm^2^). Each dot represents the average scar area of an individual heart derived from 4 sections. Increased fibrotic area was noted in mutant hearts compared to WT controls (**J**). Statistical significance was calculated using one-way ANOVA multiple comparison test. P values: ** = 0.0039, ***=0.0005

*hmmr* and *foxm1* were selected as candidates for further analysis since they are active during mammalian heart development [49-51] and mitosis [52-54]. *Hmmr* (hyaluronan-mediated motility receptor) is implicated in cell proliferation via Erk1/2 phosphorylation [55] and spindle fiber orientation [52]. It is expressed in the cardiac jelly during heart development [50], and *hmmr* activates zebrafish epicardial cells during cardiac regeneration [48]. *foxm1* is a transcription factor involved in cell cycle progression that promotes the expression of downstream genes necessary for mitotic completion [42, 53, 54, 56]. Expression of *Foxm1* is detected in the compact myocardium, trabeculated myocardium, and endothelial cells during murine heart development [49, 51] and *Foxm1* is required for cardiomyocyte proliferation [49, 51, 57] in the embryonic mouse heart. We confirmed *Foxm1* and *Hmmr* are expressed in cycling cardiomyocytes from single cell RNA sequencing (scRNA-seq) data of E10.5 embryonic mouse hearts (**Fig. 1D-F**). *Foxm1* and *Hmmr* expression was enriched in G_2_/M phase cardiomyocytes, suggesting that they can be used to identify actively cycling cells.

To determine if these genes were required for cardiac regeneration, *foxm1*^*sa10708/10708*^ (referred to as *foxm1*^*m/m*^) and *hmmr*^*sa12528/12528*^ (referred to as *hmmr*^*m/m*^) mutant lines from the Sanger zebrafish mutagenesis project [58] were studied. In both cases, homozygous mutants were viable as adults and appeared normal. We performed ventricle resection and collected hearts at 30dpa to determine if cardiac regeneration was impaired (**Fig. 1G-J**). Using acid fuchsin orange G (AFOG) staining, we observed significant presence of fibrotic tissue in *foxm1*^*m/m*^ (**Fig. 1H, J**) and *hmmr*^*m/m*^ (**Fig. 1I, J**) hearts at 30dpa compared to WT controls (**Fig. 1G**). This suggests that *foxm1* and *hmmr* are important for the progression of cardiac regeneration and their loss inhibits critical processes required for fibrotic resolution.

### *foxm1* is expressed in border zone cardiomyocytes and is required for proliferation after ventricular resection

Since Hmmr has previously been studied in mice and zebrafish [48, 50], we focused on *foxm1* as it is not known which cells express this transcription factor during adult zebrafish cardiac regeneration. Fluorescence *in situ* hybridization showed that *foxm1* mRNA was expressed in myocardium after injury (**Fig. 2A, B**). Using *Tg(myl7*:*GFP*) and *Tg(wt1b*:*GFP*) hearts, which label cardiomyocytes and epicardial cells, respectively, we confirmed that *foxm1* expression is present in *myl7*:GFP^+^ cardiomyocytes from 3dpa to 7dpa (**Fig. S1**). *foxm1*^+^ cardiomyocyte distance from the wound border was quantified and *foxm1*^+^ cells were within 100 μm from the injury plane (**Fig. 2C**). This data shows that *foxm1* is predominantly expressed in border zone cardiomyocytes at 3dpa.

**Figure 2:**
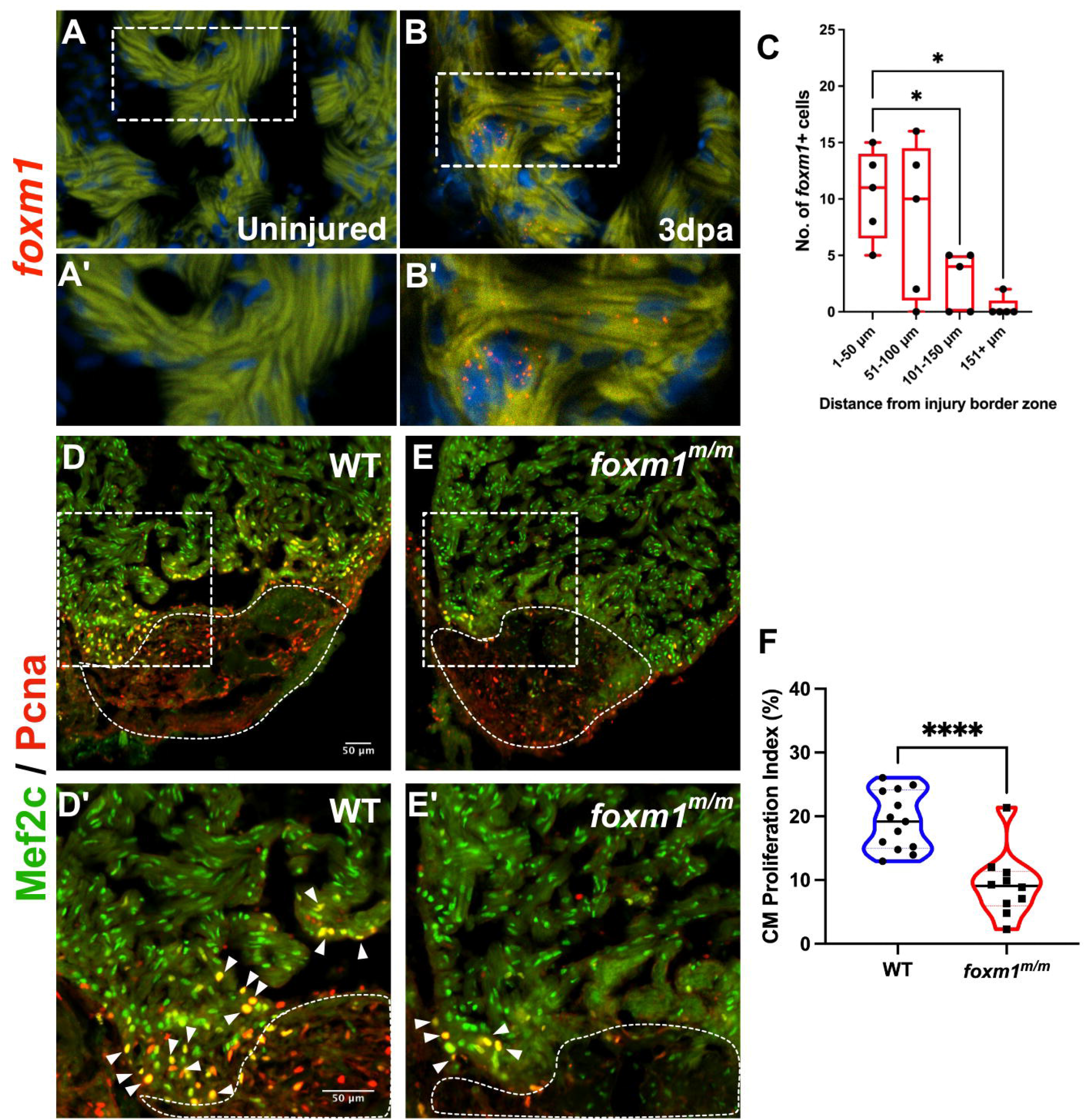
Loss of *foxm1* significantly limits cardiomyocyte proliferation after ventricular resection. Fluorescence *in situ* revealed *foxm1*(red) is not expressed in the uninjured myocardium (**A-A**’
s) but is detected in border zone cardiomyocytes after injury (**B-B**′). The myocardium is denoted by 488nm as background fluorescence. *foxm1*^+^ cardiomyocytes were localized within 100μm from the injury and expression decreased distal to the wound (**C**). Each point is representative of an individual heart (n = 5). WT (**D-E; n=13)** and *foxm1*^*m/m*^ (**F-G; n=10**) hearts at 7dpa stained for the cardiomyocyte nuclei marker Mef2c (green) and proliferation marker Pcna (red). Cardiomyocyte proliferation index shows a statistically significant decrease in *foxm1*^*m/m*^ hearts post-resection (**H**). White dotted lines represent the injury border. Each point on the violin plot represents an individual heart and these data represent 3 biological replicates. Statistical significance was calculated using one-way ANOVA multiple comparison test. P values: * < 0.05, ****= □0.0001. Scale Bars = 50μm

We next explored if *foxm1* was required for the proliferation of these cells after ventricular resection. Using WT (**Fig. 2D, D’**) and *foxm1*^*m/m*^ (**Fig. 2E, E’**) 7dpa hearts, we measured the proliferation index and observed a significant decrease in Mef2c^+^/Pcna^+^ cardiomyocytes in *foxm1* mutants (**Fig. 2F**). Together, these findings show that *foxm1* is required for cardiomyocyte proliferation and is restricted to a subset of the border zone myocardium after cardiac injury.

### *foxm1* is epistatic to *dusp6* and the Ras/MAPK pathway

We next sought to identify potential upstream pathways that regulate *foxm1* expression and activity. The EGF family ligand Neuregulin 1 (Nrg1) is expressed in the endocardium and coronary artery microvasculature in adult, mammalian hearts [59]. Nrg1 can induce cardiomyocyte division post-injury following injection of recombinant NRG1 or constitutive activation of its cognate receptor Erbb2 [60, 61]. In adult zebrafish, *nrg1* is expressed in epicardial cells from the perivascular compartment and can also stimulate cardiomyocyte proliferation after ventricular resection [30]. We hypothesized that *foxm1* expression in cardiomyocytes is localized near *nrg1*^+^ epicardial-derived cells. Double *in situ* hybridization revealed *nrg1*^+^ cells near the site of injury and were often flanked by multiple *foxm1*^+^ cardiomyocytes (**Fig. 3A, B**). The number of *foxm1*^+^ cardiomyocytes near *nrg1*^*+*^ non-myocytes was counted and a higher incidence of expression of these two genes in proximity was observed (**Fig. 3C**). As Nrg1 is known to activate the Ras/Mapk pathway and Foxm1 nuclear localization is regulated by ERK phosphorylation [62], we explored the genetic interaction between *dusp6* and *foxm1*. Given that *dusp6*^*m/m*^ hearts show increased cardiomyocyte proliferation after injury [45], we hypothesized that the *foxm1* mutation would be sufficient to block the pro-regenerative phenotype present in *dusp6* mutants. Double *dusp6:foxm1* mutant hearts showed a significant decrease in cardiomyocyte proliferation when compared to *dusp6*^*m/m*^ hearts (**Fig. 3D-H**). This demonstrates that the loss of *foxm1* is sufficient to block the pro-regenerative phenotype present in *dusp6* mutant hearts and supports an epistatic relationship where *foxm1* functions downstream of *dusp6* in heart regeneration.

**Figure 3:**
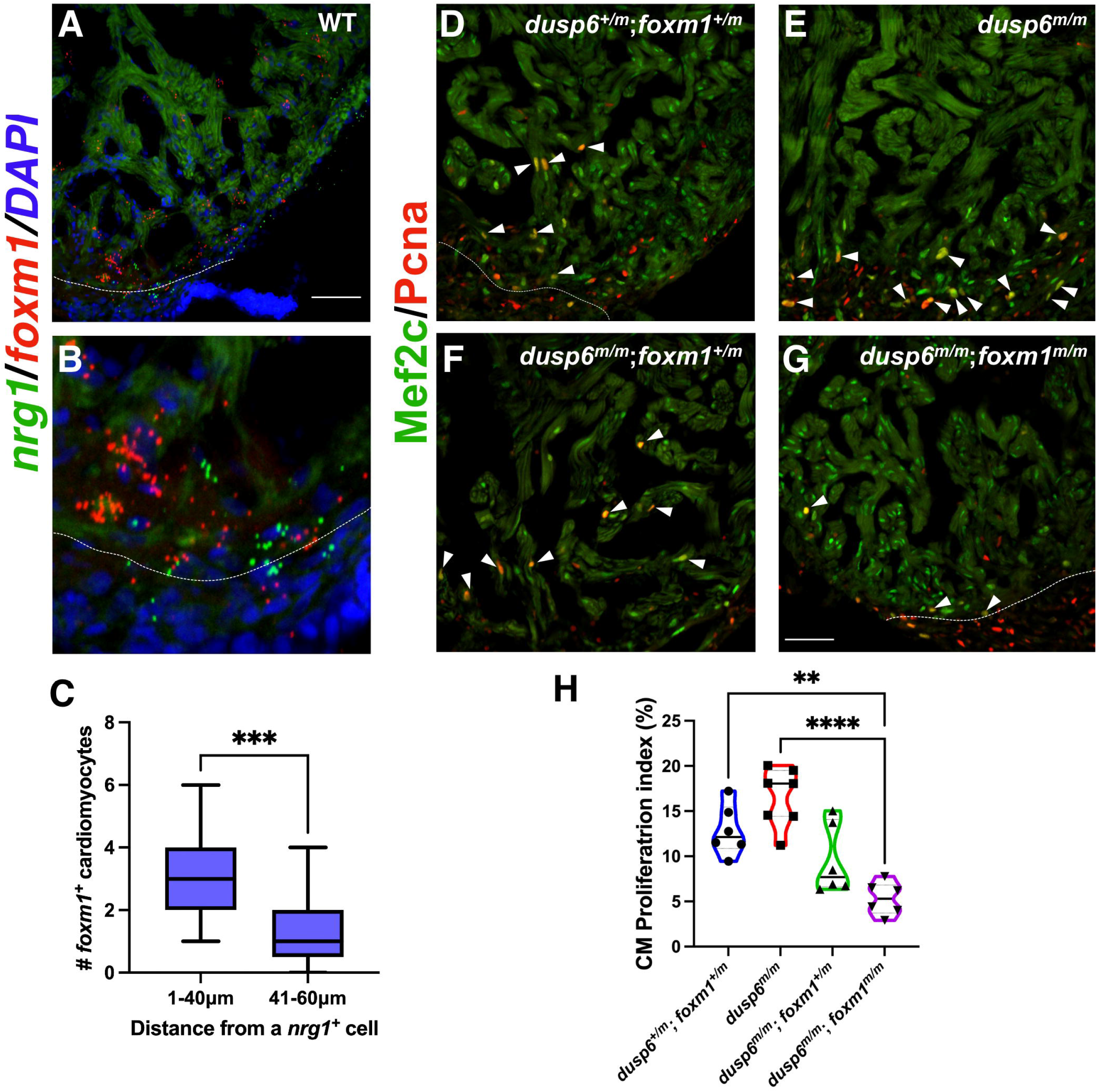
*foxm1*^+^ cardiomyocytes are localized near *nrg1*^+^ cells and is epistatic to the Mapk regulator, *dusp6*. In situ hybridization showing *nrg1* (green) and *foxm1* (red) in WT 7dpa hearts (**A**-**B**). *foxm1*^+^ cardiomyocytes were proximal to *nrg1*^+^ non-myocytes (**C**). A total of 21 *nrg1*^+^ cells were scored in 5 WT 7dpa hearts. *dusp6*^*m/*+^;*foxm1*^*m/*+^ fish were bred to generate double mutants to assess their epistatic relationship (**D-G**). *dusp6*^*m/m*^ hearts (**E; n = 7**), *dusp6*^*m/*+^;*foxm1*^*m/*+^ (**D; n = 6**) and *dusp6*^*m/m*^;*foxm1*^*m/*+^ hearts (**F; n = 6**) *dusp6*^*m/m*^;*foxm1*^*m/m*^ hearts (**G; n = 6**) were stained with Mef2c (green) and Pcna (red). Graph showing CM proliferation index (**H**). White dotted lines represent the injury border. Each point on the violin plot represents an individual heart and these data represent 2 biological replicates. Statistical significance was calculated using unpaired student’s t-test and one-way ANOVA multiple comparison test. P values: ***=0.0006 (unpaired student’s t-test: Box and Whisker plot), **=0.0022, ****<0.0001 (one-way ANOVA: Violin plot). Scale Bar = 50μm

### Cell cycle progression gene transcripts are reduced in *foxm1*^*m/m*^ hearts after injury

*Foxm1* is known to activate transcription of G_2_/M phase proliferation genes crucial for the completion of mitosis [42, 63]. Therefore, transcriptome profiling of *foxm1* mutants during heart regeneration was performed. WT and *foxm1*^*m/m*^ ventricles were collected after injury for RNA-seq, and 246 genes were significantly increased while 159 genes were decreased in *foxm1*^*m/m*^ 3dpa hearts (**Fig. 4A, and Supplementary Table 3**). We observed decreased expression of genes involved in cell cycle progression, insulin signaling, glycolysis, and AP-1 transcription factors (**Fig. 4A, and Supplementary Table 3**). Functional annotation clustering revealed that cell division and insulin signaling were decreased while immune responses were increased in *foxm1*^*m/m*^ 3dpa ventricles (**Fig. 4B, Supplementary Table 4**). qPCR validation confirmed that G_2_-phase genes (*ccnf, ccng2*, and *g2e3*) and M-phase genes (*ccnb3, cenpf*, and *prc1b*) were decreased in *foxm1*^*m/m*^ ventricles (**Fig. 4C**). In addition, genes involved in insulin signaling (*irs1, igfbp1a*, and *igfbp6b*), glycolytic genes (*pfkfb4b* and *pkmb*), and AP-1 transcription factors (*fosab, fosb*, and *jund*) were also decreased (**Fig. 4C**). These genes function in pathways known to be critical for zebrafish heart regeneration, suggesting that Foxm1 may regulate their expression after injury [64-66].

**Figure 4:**
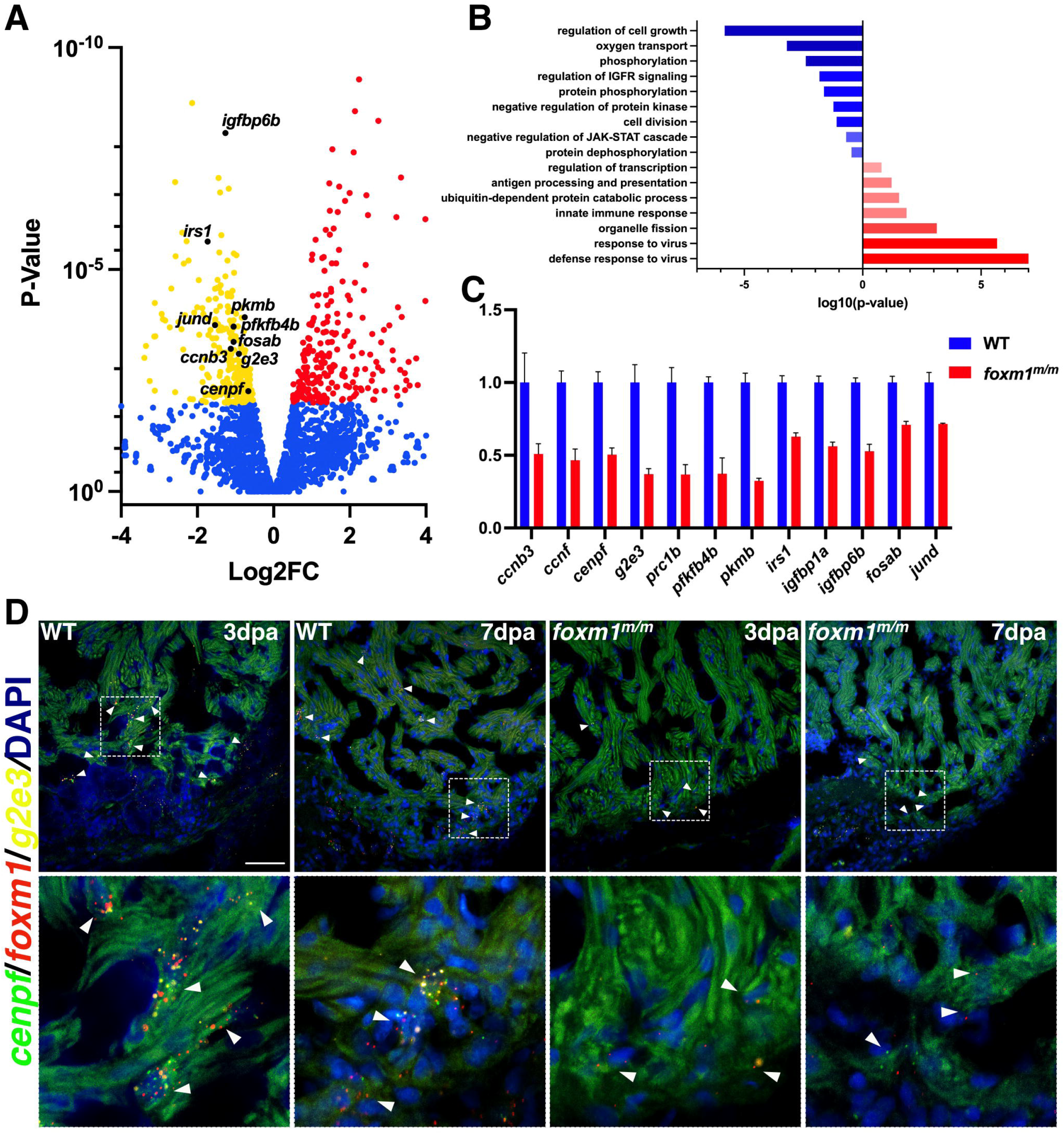
Injured *foxm1*^*m/m*^ hearts show reduced expression of G_2_ and M phase cell cycle genes. Representative Volcano plot showing WT vs *foxm1*^*m/m*^ 3dpa gene expression from RNA-seq (**A**). Genes involved in cell cycle progression, insulin signaling, glycolysis and the AP-1 transcription factor family were significantly downregulated in the *foxm1* mutant hearts. Red and yellow points indicate increase and decrease expression, respectively. Functional annotation clustering (NIH DAVID) confirmed decreased activity of cell division and insulin signaling, while innate immune response is increased (**B**). qPCR validation on a subset of the genes with data represented as fold change normalized to WT 3dpa expression + S.D. (**C**). Fluorescent *in situ* hybridization of *foxm1* and the cell cycle genes *g2e3* and *cenpf* show co-expression in a subset of border zone cardiomyocytes in WT 3dpa and 7dpa hearts (**D**) while *foxm1*^+^ *g2e3*^+^ *cenpf*^+^ cardiomyocytes were rarely detected in the *foxm1* mutants (**D**). Scale Bar = 50μm

To determine if these genes were expressed in cardiomyocytes after injury, *in situ* hybridization on WT and *foxm1*^*m/m*^ injured hearts was performed. Insulin receptor substrate 1 (*irs1*) was highly expressed in WT myocardium but was decreased in *foxm1*^*m/m*^ hearts at 3dpa (**Fig. S2A, B**). Also, protein regulator of cytokinesis 1b (*prc1b*) was expressed in border zone cardiomyocytes at 3dpa but its expression was decreased in *foxm1*^*m/m*^ hearts (**Fig. S2C, D**). Further, triple fluorescence *in situ* hybridization using *foxm1, g2e3* (G_2_ phase), and *cenpf* (M phase) revealed co-expression in cardiomyocytes near the injury border zone at 3dpa and 7dpa (**Fig. 4D**). These *foxm1, cenpf* and *g2e3* triple-positive cardiomyocytes were not detected in *foxm1*^*m/m*^ hearts, suggesting that mutant cardiomyocytes are not progressing into the latter phases of the cell cycle (**Fig. 4D**). Moreover, the triple-positive cardiomyocytes were present within 100 µm from the injury border (**Fig. S2E**), suggesting *foxm1* is required for border zone cells to cycle through G_2_/M phases and identifies a proliferative sub-population that activates post-injury. Taken together, these data suggest that *foxm1* regulates multiple pathways required for cardiomyocyte proliferation and that the loss of this transcription factor prevents the proliferative switch, thereby impairing cardiac regeneration.

### *cenpf* mutants fail to regenerate due to increased cardiomyocyte binucleation

We next investigated the importance of *cenpf*, a known Foxm1 target gene. *Cenpf* plays a role in sister chromatid separation during mitosis [43, 67, 68] and is required for normal heart development [69, 70]. In mice, *Cenpf* mutant hearts possess thinner ventricular walls, reduced heart size [69], and decreased cardiomyocyte proliferation between P2-P5 [70]; however, its role in cardiac regeneration has not been characterized. We hypothesized that *cenpf* mutant hearts would display a similar phenotype to *foxm1* mutants after ventricular resection. The *cenpf* ^*sa12296/sa12296*^ line (referred to as *cenpf* ^*m/m*^) contains a premature stop codon within Exon 6 [58]. This mutation is predicted to cause the loss of the entire C-terminal region and prevent Cenpf from interacting with spindle fibers during mitosis. *cenpf* ^*m/m*^ fish survived to adulthood with no cardiac abnormalities. After ventricular resection, *cenpf* ^*m/m*^ hearts retained large collagen/fibrin scars compared to WT controls at 30dpa (**Fig. 5A-C**). Furthermore, fibrotic scars were still evident in *cenpf*^*m/m*^ hearts as late as 60dpa compared to controls (**Fig. S3**). We next determined if cardiomyocyte proliferation was affected by the loss of *cenpf*. In contrast to *foxm1*^*m/m*^ hearts, no significant difference in the number of Pcna^+^ cardiomyocytes were detected in *cenpf*^*m/m*^ hearts after injury (**Fig. 5D-F**). However, fluorescence *in situ* hybridization revealed a decrease of *prc1b* expression in *myl7:*GFP^+^ cardiomyocytes in *cenpf*^*m/m*^ hearts (**Fig. 5H**) compared to controls (**Fig. 5G**). With prolonged fibrosis and decreased expression of the cytokinesis marker *prc1b* in *cenpf* mutants post-injury, we postulated that cytokinetic dysfunction occurred in the regenerating cardiomyocytes, thereby preventing the formation of healthy myocardium in the mutants.

**Figure 5:**
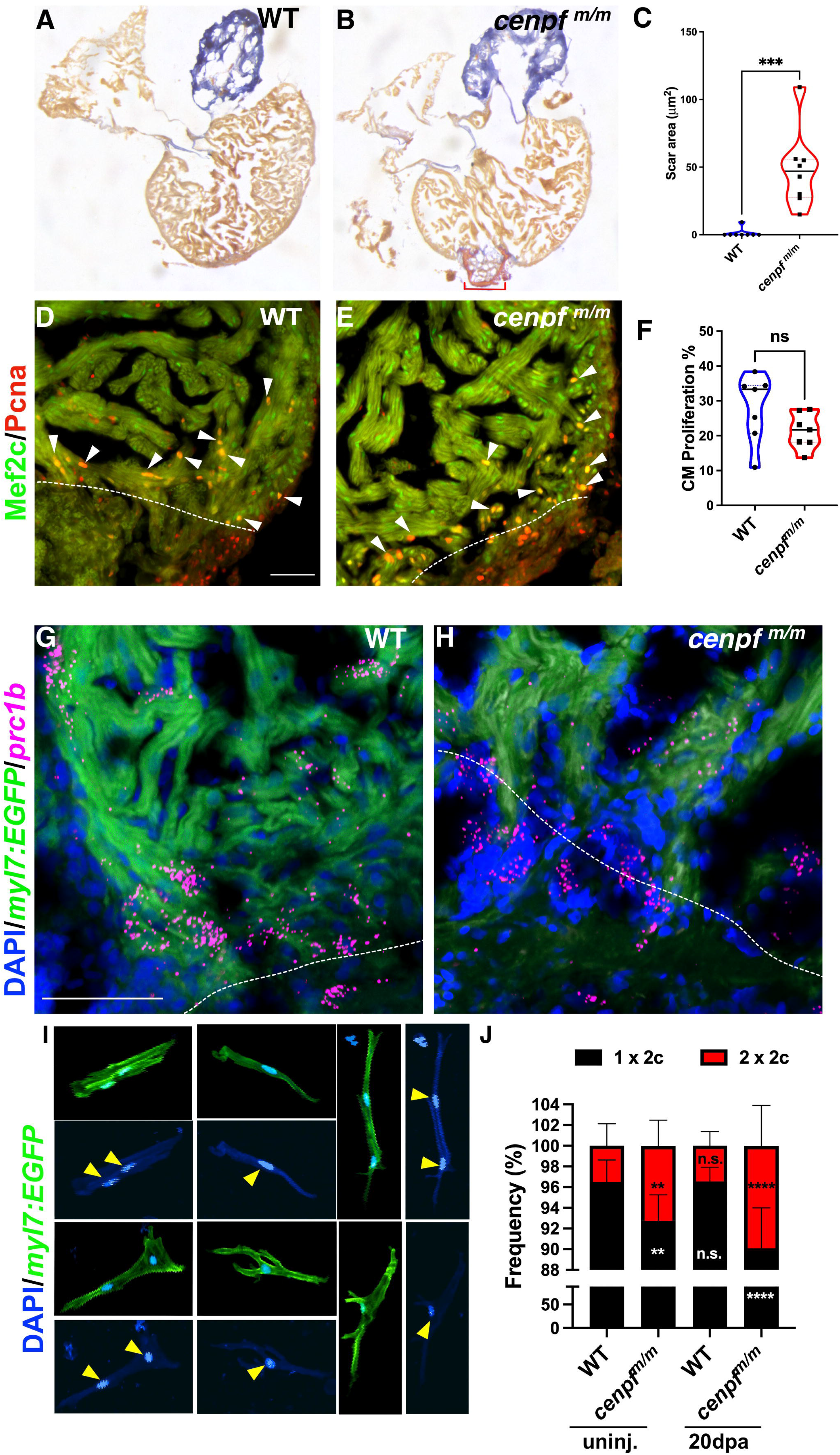
Proportion of binucleated cardiomyocytes were increased in *cenpf*^*m/m*^ hearts. WT (**A; n = 8**) and *cenpf*^*m/m*^ (**B; n = 8**) 30dpa hearts stained with AFOG to label fibrotic tissue. Scar area was calculated as pixel value and converted into microns squared (μm^2^). Each dot represents the average scar area of an individual heart derived from 4 sections (**C**). AFOG staining was used to label fibrin (red), collagen (blue), and muscle (orange-brown). WT (**D; n =7**) and *cenpf*^*m/m*^ (**E; n = 7**) hearts showed no significant difference in cardiomyocyte proliferation index (**F**). White dotted lines represent the injury border. Fluorescent *in situ* showed *prc1b* (magenta) was present in GFP^+^ cardiomyocytes in WT hearts (**G**) but *prc1b*^+^ cardiomyocytes were decreased in *cenpf*^*m/m*^ 7dpa hearts (**H**). Individual cardiomyocytes were isolated from uninjured and 20dpa hearts extracted from Tg(*myl7:EGF)* WT and *cenpf*^*m/m*^ zebrafish (**I, J**) to count mononucleated vs binucleated cardiomyocytes. Representative images of mononucleated and binucleated cardiomyocytes are shown (**I**). The frequency (%) of WT uninjured (n = 11 hearts; 1945 CMs; binucleation = 3.52%), *cenpf*^*m/m*^ uninjured (n = 11; 3204 CMs; binucleation = 7.24%), WT 20dpa (n = 11; 1858 CMs; binucleation =3.45%), and *cenpf*^*m/m*^ 20dpa (n = 12; 3072 CMs; binucleation = 9.90%) mononucleated (1 × 2c; black) vs binucleated (2 × 2c; red) cardiomyocytes were determined. Statistical analysis for AFOG and proliferation index was calculated using the unpaired student’s t-test while mononucleated vs binucleated counts used two-way ANOVA with Dunnett’s multiple comparisons test. P values: ** ≤ 0.0046, *** = 0.0004, **** □ 0.0001. Scale Bar = 50μm

During mammalian development, the heart primarily contains mononucleated, diploid cardiomyocytes (1 × 2c) but after P7 they either become mononucleated polyploid or binucleated [23, 24, 27, 28]. Unlike mammals, the majority of adult zebrafish cardiomyocytes remain mononucleated with less than 5% of cardiomyocytes being binucleated [46, 71, 72]. Cytokinesis occurs after nuclei division and is characterized by cleavage furrow constriction and abscission [73]. Binucleation is caused by cytokinetic failure and is a hallmark of cardiomyocyte maturation in mammals [74-76]. Since Cenpf is known to associate to the kinetochore and spindle fibers prior to cytokinesis, we investigated if the loss of *cenpf* increased cardiomyocyte binucleation after cardiac injury. We generated a *Tg*(*myl7*:*GFP*); *cenpf*^*m/m*^ zebrafish to count nuclei within individual, GFP^+^ cardiomyocytes. Mononucleated diploid and binucleated cardiomyocytes were detected in these isolated cells (**Fig. 5I**). Binucleated cells comprised 3.52% of the cardiomyocytes isolated from uninjured hearts and remained constant in WT 20dpa hearts at 3.45%, indicating that no significant increase occurred post-injury (**Fig. 5J**). However, the percentage of binucleated cardiomyocytes increased by 3.68% in uninjured *cenpf*^*m/m*^ hearts, which was double the amount detected in WT controls (**Fig. 5J)**. And after ventricular resection, the percentage of binucleated cardiomyocytes increased by 6.38% at 20dpa in *cenpf* mutant hearts compared to controls (**Fig. 5J**). Therefore, the loss of *cenpf* caused mitotic dysregulation during heart regeneration, and because fewer cardiomyocytes divided, fibrotic scar tissue persisted.

## Discussion

In this study, we demonstrate transcriptional regulation of late-stage, cell cycle progression genes is essential for post-injury cardiomyocyte proliferation during heart regeneration. Foxm1 is necessary for cardiomyocyte proliferation through the induction of G_2-_ and M-phase genes after injury (**Fig. 6**). Moreover, cell cycle progression genes, such as *cenpf*, are required for successful cardiomyocyte division after cardiac injury (**Fig. 6**).

Multiple factors can stimulate cardiomyocyte proliferation in adult zebrafish and neonatal mouse hearts after cardiac injury. Secreted ligands, such as *nrg1, bmp2b, tgf-β1, pdgfbb, igf2b*, and *fgf17b*, are produced by non-myocytes during cardiac injury and stimulate cardiomyocyte proliferation [30, 32, 33, 60, 77-83]. Recent studies have also highlighted the role of transcription factors that regulate cardiomyocyte proliferation in adult zebrafish and neonatal mice post-injury. These include the AP-1 family transcription factors Fos and Jun [64, 84-86]. Fos and Jun dimerize and bind to AP-1 motifs to induce cell proliferation. It was recently shown that AP-1 motifs in chromatin are readily accessible in regenerating cardiomyocytes [64]. Several groups report the AP-1 factors *junbb, fosl1a*, and *fosab* are expressed in border zone cardiomyocytes and are required for dedifferentiation and proliferation [64, 84, 86]. In the post-MI adult mouse heart, *Jun* or *Fosl1* are not expressed in the border zone cardiomyocytes [85]. This highlights the importance of inducing dedifferentiation and proliferation by AP-1 transcription factors in mature cardiomyocytes to stimulate them into a regenerative state.

Foxm1 was previously shown to directly bind to DNA via forkhead recognition elements in promoters to regulate gene expression [42]. More recent studies show Foxm1 can bind to DNA by forming a complex with the transcription factor MuvB or through interactions with non-consensus domains to promote transcription [87-90]. Foxm1 activates transcription of G_2_/M-phase cell cycle genes, DNA damage response genes, and cell migration genes [42, 53, 54, 63, 87-91]. We observed decreased expression of G_2_/M-phase cell cycle genes in *foxm1* mutant hearts, but we also detected decreased expression of AP-1 transcription factors and glycolysis genes. Previous studies suggest that FoxM1 transcriptionally activates *JNK1*, a kinase that phosphorylates the AP-1 factor *c-Jun*, to promote cell migration and invasion in human U2OS osteosarcoma cells [39]. In addition, an AP-1 motif was detected within the FoxM1 promotor in human cancer cells and c-Jun binds to the promotor region [92]. We noted decreased transcripts for *fosab, fosb*, and *jund* in *foxm1*^*m/m*^ 3dpa hearts, which suggests a positive feedback loop between AP-1 and Foxm1 may exist to promote cardiomyocyte dedifferentiation and proliferation.

The metabolic switch from oxidative phosphorylation to aerobic glycolysis (the Warburg effect [93]) has been well studied in cardiac biology. Mature cardiomyocytes are dependent on oxidative phosphorylation to generate the energy required for contraction [18], while embryonic and fetal cardiomyocytes primarily use glycolysis [94, 95]. This metabolic change occurs in border zone cardiomyocytes in both adult zebrafish and neonatal mice as blocking glycolysis in these embryonic-appearing cardiomyocytes significantly reduce their proliferation [65, 66]. *FOXM1* is known to bind to the promoter regions of *LDHA*, an enzyme that converts pyruvate into lactate [96]. FoxM1 also binds to the promoters of glycolysis rate-limiting genes *GLUT1*, a transmembrane protein that transports glucose into the cell, and *HK2*, an enzyme that converts glucose into glucose-6-phosphate [97] in human cancer lines. In *foxm1*^m/m^ hearts, we found the rate-limiting glycolysis genes *pfkfb4b* and *pkmb* were decreased. Therefore, zebrafish Foxm1 may play a role in promoting transcription of these glycolysis genes in the border zone cardiomyocytes to enhance the switch from oxidative phosphorylation toward glycolysis to sustain proliferation.

Recent studies show that mutations within genes associated with the cell cycle can increase the number of mononucleated polyploid and binucleated cardiomyocytes, which attenuate heart regeneration [46, 71, 98, 99]. *ect2* encodes a protein required for cleavage furrow abscission during cytokinesis was shown to reduce cardiomyocyte binucleation when overexpressed in neonatal rat cardiomyocytes [76]. In contrast, inactivation of *etc2* via knockout or ectopic expression of a dominant-negative isoform caused both binucleation and polyploidization in neonatal rat and adult zebrafish cardiomyocytes [46, 76]. Cenpf is a facilitator of sister chromatid separation as it associates to the kinetochores from prometaphase to early anaphase [43, 44] and silencing of Cenpf disrupts chromosome alignment and can cause cytokinetic failure [41]. In this study we observed an accumulation of binucleated cardiomyocytes after ventricular resection in *cenpf* mutant hearts. This demonstrates that *cenpf*^*m/m*^ cardiomyocytes display disrupted cytokinesis generating a ploidy defect consistent with impaired regeneration.

Overall, we demonstrate that *foxm1* and *cenpf* are expressed in a specific sub-population of regenerating cardiomyocytes at the injury border. Disrupting the Foxm1-Cenpf axis reduced cardiomyocyte proliferation and mitosis via decreased G_2_/M cell cycle gene expression. These results illustrate the importance of a transcription factor in cardiomyocyte proliferation, and these studies could assist in designing strategies to improve regeneration in the adult mammalian heart [100].

## Acknowledgements

This research was supported by funding from the National Institutes of Health (NIH) (R01HL142788 and R01HL156398 to MT; R01HL157879 and R01HL159805 to PVB; 1F31HL149148 and T32EB001026 to DAZ). CLC Genomics Workbench software licensed through the Molecular Biology Information Service of the Health Sciences Library System, University of Pittsburgh was used for data analysis. This research was supported in part by the University of Pittsburgh Center for Research Computing through the resources provided. We are grateful to members of the Tsang lab, Donghun Shin, Andrew Duncan, Bernhard Kuhn and Neil Hukriede for reading the manuscript and experimental suggestions.

## Data Resource

The RNA-seq data files are available under the accession number: GSE201139

## Materials and Methods

### Zebrafish maintenance and ventricular amputation

All zebrafish experiments and protocols were performed according to protocols approved by the Institutional Animal Care and Use Committee (IACUC) at the University of Pittsburgh in agreement with NIH guidelines. Adult (6-to 18-month-old) wild-type AB*, Tg(*myl7:GFP*)^twu34Tg^ [101], Tg(*wt1b:EGFP*)^*li1*^ [102], mutant *dusp6*^*pt30a*^ [45], mutant *foxm1*^*sa10708*^, mutant *hmmr*^*sa12528*^, and mutant *cenpf*^*sa12296*^. *foxm1*^*sa10708*^, *hmmr*^*sa12528*^, and *cenpf*^*sa12296*^ were acquired from the Zebrafish International Resource Center (ZIRC) [58].

DNA was isolated using Proteinase K (10mg/mL) denaturation following adult tail fin clips for genotyping assays. PCR genotyping for *dusp6*^*pt30a pt30a*^, *foxm1*^*sa10708/sa10708*^, *cenpf*^*sa12296/sa12296*^, and *hmmr*^*sa12528/12528*^ (**Supplementary Table 5**) was performed and products were digested with the restriction enzymes Cla1, Sal1, Spe1, or Age1, respectively. TaqMan SNP assays (Thermo Fisher Scientific) using probes for *cenpf* (AN33F96, Thermo Fisher Scientific) and *foxm1* (AN2XMN9, Thermo Fisher Scientific) were also used to identify mutant SNPs.

Ventricular amputation was performed as previously described [5] with approximately 20% of the ventricle apex being resected. Zebrafish were returned to the aquarium for standard feeding and husbandry before hearts were extracted for DNA, RNA, or histology at specific time points.

### RNA extraction, cDNA, synthesis, and quantitative PCR (qPCR)

Total RNA was extracted from uninjured and injured (3 and 7dpa) hearts using Trizol (Invitrogen) and was purified with the RNeasy Micro Kit (Qiagen). A minimum of eight hearts were pooled together for each condition. 1μg of RNA was reverse transcribed into cDNA using SuperScript reverse transcriptase (Invitrogen) with random hexamers. qPCR was performed as previously described[48]. β-actin2 (*bactin2*) and RNA polymerase II subunit D (*polr2d*) were used to normalize gene expression in these experiments. qPCR primers are listed in Supplementary Table 5. At least two, independent biological replicates were performed.

### RNA-seq sample preparation and data analysis

Total RNA was extracted from uninjured and 3dpa ventricles from WT, *dusp6*^*pt30a*^*/*^*pt30a*^, and *foxm1*^*sa10708/sa10708*^ adult zebrafish. A minimum of eight hearts were pooled together for each condition and RNA was isolated as previously described. For RNA-seq, 0.1-0.5 μg for each condition and sent RNA to the Genomics Research Core at the University of Pittsburgh, and to Tufts University for library preparation and sequencing. The raw sequence reads were processed and mapped to the Zebrafish Reference Genome GRCz11 using CLC Genomics Workbench 20. Differentially expressed genes were calculated within the CLC software package and classified based on FDR of ≤ 0.01 and LOG2FC ≥ 0.59 for uninjured vs 3dpa experiment. For the *foxm1* mutant heart RNA-seq study, differential expression was based on FDR ≤0.05 and LOG2FC ≥ 0.59.

### scRNA-seq data analysis

The mouse single cell data was generated previously (PMID: 31142541) and downloaded from GEO under the accession number: GSE122403. Isolated cardiomyocytes from left ventricles were used for the analysis including cell cycle phase annotation and gene expression analysis in Seurat V3 following a procedure as described previously (PMID: 31142541).

### AFOG Fibrosis Assay

For fibrosis assays, uninjured and injured hearts were collected in cold 1X PBS and fixed in 4% PFA in PBS for either 2 hours at room temperature (RT) or overnight at 4ºC. Hearts were transferred into a sucrose gradient (10-20-30% sucrose in 1X PBS; ∼1 hr for each solution) and stored in 30% sucrose in 1x PBS at 4ºC overnight. Hearts were embedded in Surgipath Cryo-Gel (Leica, 39475237) the following day, and samples were sectioned on a cryostat at 14 µm and were dried overnight at RT. For fibrosis staining, we performed Acid Fuchsin Orange G (AFOG) staining as previously described [5]. Images were taken with a Leica MZ16 microscope and Q Imaging Retiga 1300 camera. Fibrotic areas were measured using FIJI. For each heart, the average of the sum of the scar area (μm^2^) was calculated from the four largest central sections.

### Immunofluorescent Staining

For immunofluorescent (IF) staining, slides washed in PBS-T (2x, 5 min), deionized water (2x, 5min), and were then permeabilized using 4% hydrogen peroxide in 100% methanol for 1 hour at RT. After permeabilizing the sections, they were washed in deionized water (2x, 5min) then PBS-T (2x, 5 min), and then incubated in IF buffer (200uL sheep serum, 100 uL DMSO, 20 uL TritonX-100, and 9680 uL of PBS for 10mL) for a minimum of 30 min at RT. Primary antibodies used for IF staining were rabbit polyclonal anti-Mef2c (Santa Cruz, sc-313; 1:500) and mouse monoclonal anti-Pcna (Sigma, P8825; 1:3000). Secondary antibodies were Alexa Fluor 488 goat anti-rabbit IgG peroxidase conjugate (Invitrogen, A11008; 1:1000) and Alexa Fluor 594 goat anti-mouse IgG (H+L) (Invitrogen, A11005; 1:1000). Incubation with primary and secondary antibodies was done overnight at 4ºC. In between each incubation, slides were washed in PBS-T (6x, 5 min) at RT to remove unbound antibodies. Lastly, slides were treated with DAPI (1:500, 3 min RT, ThermoFisher Scientific, D1306), washed briefly in PBS (2x), and then sealed using Aqua-PolyMount (Polysciences, Inc, #18606-20). Images were taken on a Zeiss 700 confocal microscope using a 20X objective. Image analysis was performed using ImageJ Fiji (NIH).

### Cardiomyocyte Proliferation Index

After immunofluorescent staining of heart sections with Mef2c and Pcna, the cardiomyocyte proliferation index (%) was calculated from the number of Mef2c^+^/Pcna^+^ divided by the total number of Mef2c+ cells. A total of four sections were counted per individual heart and a minimum of two biological replicates used in each experiment.

### RNAscope

RNAscope [Advanced Cell Diagnostics (ACD)] was performed on uninjured and injured (3, 7, and 14 dpa) hearts isolated from AB*, Tg(*myl7:GFP*)^twu34Tg^, Tg(*wt1b:EGFP*)^*li1*^, and mutant *foxm1*^*sa10708*^ adult zebrafish. Hearts were fixed overnight at 4°C in 4% paraformaldehyde (PFA), transferred into a sucrose gradient (10-20-30%) the following day at 4°C before cryopreservation overnight. Tissue was embedded in Surgipath Cryo-Gel and sectioned at 14 µm. RNAscope probe hybridization, amplification and immunostaining were performed following the protocol provided in the RNAscope Multiplex Fluorescent Reagent Kit v2 user manual (ACD). ACD designed all the probes used in this study and they are listed in Supplementary Data Table 6. For some sections following the final wash step of the RNAscope probe hybridization protocol, immunofluorescent staining was performed to better visualize endogenous GFP from the transgenic lines. We started our IF protocol by treating the slides with IF and then performing the rest of the protocol as normal. Primary antibodies for immunostaining were chicken anti-GFP (1:1000; Aves Labs, GFP-1020). Secondary antibodies for immunostaining were fluorescein goat anti-chicken 488 (1:1000; Aves Labs, F-1005). Slides were treated with DAPI (1:500, 3 min RT). Images were taken on a Zeiss 700 confocal microscope using a 40X water objective. Image analysis was performed using ImageJ Fiji (NIH).

### Cardiomyocyte dissociation

We performed ventricular resections on Tg(*myl7:GFP*)^twu34Tg^ and Tg(*myl7:GFP*)^twu34Tg^:*cenpf*^*m/m*^ adult zebrafish and collected hearts at 20dpa. For a crude dissociation, we used the assay previously described by González-Rosa *et al*. [46]. Individual hearts were extracted and placed in ice-cold dissection buffer (0.3% bovine serum albumin and 20mM glucose in 1X PBS) before removing the bulbous arteriosus and atrium from the ventricle. The ventricles were cut into smaller pieces and washed twice in fresh dissection buffer for 5 min at RT. Ventricles were then treated with digestion solution (20mM glucose 10mM BDM into 1X Trypsin-EDTA (ThermoFisher Scientific, Cat# 2520056)) for 15 min on ice and with gentle agitation. Digestion solution was then removed and ventricles were washed 3 times with dissection buffer with BDM (0.3% bovine serum albumin, 20mM glucose, and 10mM BDM in 1X PBS) at RT. Following these washes, ventricles were treated in a collagenase digestion solution (20mM glucose and 10mM BDM in 100% Accumax solution (EMD Millipore, SCR006) for 45 min at RT and tubes were gently vortexed every 10 min. Ventricles were pipetted in the collagenase digestion solution until ventricle pieces were no longer visible and cells were pelleted on a tabletop centrifuge @ 400 x g for 5 min. Collagenase solution was removed without disturbing the pellet, 4% PFA in 1X PBS was added to the pellet, and cells were fixed for 1 hr at RT. Cells were centrifugated @ 400 x g for 5 min, and the pellet was resuspended in 1X PBS. Single cell suspension was spread onto SuperFrost Plus microscope slides outlined with a hydrophobic barrier pen and cells were allowed to flatten O.N.

### Binucleation Counts

For fixed single cell suspension, slides were permeabilized in 0.5% Triton X-100 in 1X PBS for 10 min at RT. Slides were gently washed 3 times using 1X PBS. Slides were blocked in blocking buffer (5% goat serum and 0.1% Tween 20 in 1X PBS) for 30 min at RT. Chicken anti-GFP was used as primary antibody (1:1000; Aves Labs, GFP-1020) and slides were incubated at 4ºC O.N. Slides were washed in 0.5% NP40 in 1X PBS (3x, 5 min each, RT). Fluorescein goat anti-chicken antibody was used as secondary antibody (1:200; Aves Labs, F-1005) and slides were incubated for 2 hours at RT. Slides were washed in 0.5% NP40 in 1X PBS (3x, 5 min each, RT). All slides were treated with DAPI (1:500) for 10 minutes at RT and were immediately washed in 1X PBS (2x, 5 min each, RT) before being sealed using Aqua-PolyMount. Images were taken on a Zeiss 700 confocal microscope using a 20X objective and ImageJ FIJI (NIH). GFP^+^ binucleated cardiomyocytes were manually counted along with all other GFP^+^ mononucleated cardiomyocytes and the mean percentage + S.D. of mononucleated and binucleated cardiomyocytes was represented in a stacked bar graph.

### Statistical Analysis

Statistical analyses were determined with GraphPad Prism version 9.3. Statistical significance was analyzed by unpaired Student’s t-test, one-way ANOVA, or two-way ANOVA. Data are shown as mean ± S.D or mean ± S.E.M. p<0.05 was considered significant.

## Figure Legends

**Figure 6: Foxm1 and Cenpf expression is required in border zone cardiomyocytes to permit cell cycle progression and prevent mitotic dysregulation.**In post-injury hearts, a sub-population of border zone cardiomyocytes express *foxm1*. Foxm1 transcriptionally activates cell cycle genes such as *cenpf, g2e3*, and *prc1b* to promote regeneration of the lost myocardium. In the absence of *foxm1*, G_2_/M phase genes are not expressed in sufficient levels and border zone cardiomyocytes fail to proliferate allowing fibrotic tissue to persist. In *cenpf*^*m/m*^ hearts, *foxm1* expression is still present and border zone cardiomyocytes initiate mitosis; however, the absence of *cenpf* suppresses mitosis and produces binucleated cardiomyocytes post-injury. The increase in binucleated cardiomyocytes prevents regeneration.

## Figure Supplement Legends

**Figure S1. *foxm1* is predominantly expressed in the border zone myocardium after ventricular resection**. Fluorescent *in situ* was used in Tg(*myl:GFP*) and Tg(*wt1b*:*GFP*) hearts to determine the spatial expression of *foxm1. foxm1* is not expressed in the uninjured heart (**A; n = 3**) but its expression is detected in GFP^+^ cardiomyocytes at 3dpa (**B; n = 3**) and 7dpa (**C; n = 3**) in Tg(*myl7:GFP*). In contrast, *foxm1* was not expressed in GFP^+^ epicardial cells in Tg(*wt1b*:*GFP*) 3dpa hearts (**D**). The number of *foxm1*^+^ GFP^+^ cardiomyocytes increased after injury (**E**). Statistical significance was calculated by one-way ANOVA with Dunnett’s multiple comparison test as either the mean + S.D as a bar graph or represented in a violin plot. P value: ** = 0.0077, *** = 0.0002. Scale Bar = 50μm

**Figure S2. *irs1* and prc1b are decreased in *foxm1***^***m/m***^ **hearts**. Fluorescent *in situ* was used to determine the spatial expression of *irs1* and *prc1b. irs1* is highly expressed in uninjured and 3dpa WT hearts in the myocardium (**A**) **B**). In foxm1 mutants, *irs1* is slightly decreased in *foxm1*^*m/m*^ uninjured (**C**) and 3dpa (**D**) myocardium which suggests insulin-mediated activity may be affected. *prc1b* was not detected in the WT uninjured (**E**) and *foxm1*^*m/m*^ uninjured (**G**) heart. *prc1b* expression increases in the myocardium and clot in WT 3dpa hearts (**F**), but *foxm1*^*m/m*^ 3dpa hearts predominantly express *prc1b* in the clot. Triple positive cardiomyocytes were counted, and this subset is typically found within 100μm from the clot (**E**). Triple positive cardiomyocytes were counted in total from 5 hearts. White dotted lines represent the injury border. Scale Bar = 50μm

**Figure S3. *cenpf***^***m/m***^ **60dpa hearts retain scar tissue which suggests a more permanent fibrotic phenotype**. WT (**A; n = 9**) and *cenpf*^*m/m*^ (**B; n = 7**) 60dpa were stained with AFOG solution and the scar area (μm^2^) was measured. *cenpf*^*m/m*^ hearts still retain a collagen scar at the injury site that is significantly larger than WT controls (**C**). Statistical significance was calculated using the unpaired student’s t-test and represented using mean + S.D. P value: * = 0.0119.

## References

1. Becker, R.O., S. Chapin, and R. Sherry, Regeneration of the ventricular myocardium in amphibians. Nature, 1974. 248(5444): p. 145–7.

2. Oberpriller, J.O. and J.C. Oberpriller, Response of the adult newt ventricle to injury. J Exp Zool, 1974. 187(2): p. 249–53.

3. Chablais, F., et al., The zebrafish heart regenerates after cryoinjury-induced myocardial infarction. BMC Dev Biol, 2011. 11: p. 21.

4. Gonzalez-Rosa, J.M., et al., Extensive scar formation and regression during heart regeneration after cryoinjury in zebrafish. Development, 2011. 138(9): p. 1663–74.

5. Poss, K.D., L.G. Wilson, and M.T. Keating, Heart regeneration in zebrafish. Science, 2002. 298(5601): p. 2188–90.

6. Raya, A., et al., Activation of Notch signaling pathway precedes heart regeneration in zebrafish. Proc Natl Acad Sci U S A, 2003. 100 Suppl 1(Suppl 1): p. 11889–95.

7. Porrello, E.R., et al., Transient regenerative potential of the neonatal mouse heart. Science, 2011. 331(6020): p. 1078–80.

8. Ito, K., et al., Differential reparative phenotypes between zebrafish and medaka after cardiac injury. Dev Dyn, 2014. 243(9): p. 1106–15.

9. Rumyantsev, P.P., Interrelations of the proliferation and differentiation processes during cardiact myogenesis and regeneration. Int Rev Cytol, 1977. 51: p. 186–273.

10. Agarkova, I., et al., A novel marker for vertebrate embryonic heart, the EH-myomesin isoform. J Biol Chem, 2000. 275(14): p. 10256–64.

11. Guo, Y. and W.T. Pu, Cardiomyocyte Maturation: New Phase in Development. Circ Res, 2020. 126(8): p. 1086–1106.

12. Padula, S.L., N. Velayutham, and K.E. Yutzey, Transcriptional Regulation of Postnatal Cardiomyocyte Maturation and Regeneration. Int J Mol Sci, 2021. 22(6).

13. England, J. and S. Loughna, Heavy and light roles: myosin in the morphogenesis of the heart. Cell Mol Life Sci, 2013. 70(7): p. 1221–39.

14. Kubalak, S.W., et al., Chamber specification of atrial myosin light chain-2 expression precedes septation during murine cardiogenesis. J Biol Chem, 1994. 269(24): p. 16961–70.

15. Lyons, G.E., et al., Developmental regulation of myosin gene expression in mouse cardiac muscle. J Cell Biol, 1990. 111(6 Pt 1): p. 2427–36.

16. O’Brien, T.X., K.J. Lee, and K.R. Chien, Positional specification of ventricular myosin light chain 2 expression in the primitive murine heart tube. Proc Natl Acad Sci U S A, 1993. 90(11): p. 5157–61.

17. Siedner, S., et al., Developmental changes in contractility and sarcomeric proteins from the early embryonic to the adult stage in the mouse heart. J Physiol, 2003. 548(Pt 2): p. 493–505.

18. Sartiani, L., et al., Developmental changes in cardiomyocytes differentiated from human embryonic stem cells: a molecular and electrophysiological approach. Stem Cells, 2007. 25(5): p. 1136–44.

19. Papanicolaou, K.N., et al., Mitofusins 1 and 2 are essential for postnatal metabolic remodeling in heart. Circ Res, 2012. 111(8): p. 1012–26.

20. Fritz, H.L., I.W. Smoak, and S. Branch, Hexokinase I expression and activity in embryonic mouse heart during early and late organogenesis. Histochem Cell Biol, 1999. 112(5): p. 359–65.

21. Dai, D.F., et al., Mitochondrial Maturation in Human Pluripotent Stem Cell Derived Cardiomyocytes. Stem Cells Int, 2017. 2017: p. 5153625.

22. Kang, M.J., et al., Cyclins and cyclin dependent kinases during cardiac development. Mol Cells, 1997. 7(3): p. 360–6.

23. Soonpaa, M.H., et al., Cardiomyocyte DNA synthesis and binucleation during murine development. Am J Physiol, 1996. 271(5 Pt 2): p. H2183–9.

24. Walsh, S., et al., Cardiomyocyte cell cycle control and growth estimation in vivo--an analysis based on cardiomyocyte nuclei. Cardiovasc Res, 2010. 86(3): p. 365–73.

25. Ikenishi, A., et al., Cell cycle regulation in mouse heart during embryonic and postnatal stages. Dev Growth Differ, 2012. 54(8): p. 731–8.

26. Tane, S., et al., CDK inhibitors, p21(Cip1) and p27(Kip1), participate in cell cycle exit of mammalian cardiomyocytes. Biochem Biophys Res Commun, 2014. 443(3): p. 1105–9.

27. Bergmann, O., et al., Dynamics of Cell Generation and Turnover in the Human Heart. Cell, 2015. 161(7): p. 1566–75.

28. Mollova, M., et al., Cardiomyocyte proliferation contributes to heart growth in young humans. Proc Natl Acad Sci U S A, 2013. 110(4): p. 1446–51.

29. Wang, J., et al., The regenerative capacity of zebrafish reverses cardiac failure caused by genetic cardiomyocyte depletion. Development, 2011. 138(16): p. 3421–30.

30. Gemberling, M., et al., Nrg1 is an injury-induced cardiomyocyte mitogen for the endogenous heart regeneration program in zebrafish. Elife, 2015. 4.

31. Han, Y., et al., Vitamin D Stimulates Cardiomyocyte Proliferation and Controls Organ Size and Regeneration in Zebrafish. Dev Cell, 2019. 48(6): p. 853-863.e5.

32. Lien, C.L., et al., Gene expression analysis of zebrafish heart regeneration. PLoS Biol, 2006. 4(8): p. e260.

33. Wu, C.C., et al., Spatially Resolved Genome-wide Transcriptional Profiling Identifies BMP Signaling as Essential Regulator of Zebrafish Cardiomyocyte Regeneration. Dev Cell, 2016. 36(1): p. 36–49.

34. Kalin, T.V., et al., Increased levels of the FoxM1 transcription factor accelerate development and progression of prostate carcinomas in both TRAMP and LADY transgenic mice. Cancer Res, 2006. 66(3): p. 1712–20.

35. Karadedou, C.T., et al., FOXO3a represses VEGF expression through FOXM1-dependent and - independent mechanisms in breast cancer. Oncogene, 2012. 31(14): p. 1845–58.

36. Kim, I.M., et al., The Forkhead Box m1 transcription factor stimulates the proliferation of tumor cells during development of lung cancer. Cancer Res, 2006. 66(4): p. 2153–61.

37. Madureira, P.A., et al., The Forkhead box M1 protein regulates the transcription of the estrogen receptor alpha in breast cancer cells. J Biol Chem, 2006. 281(35): p. 25167–76.

38. Pan, H., et al., Transcription factor FoxM1 is the downstream target of c-Myc and contributes to the development of prostate cancer. World J Surg Oncol, 2018. 16(1): p. 59.

39. Wang, I.C., et al., FoxM1 regulates transcription of JNK1 to promote the G1/S transition and tumor cell invasiveness. J Biol Chem, 2008. 283(30): p. 20770–8.

40. Yoshida, Y., et al., The forkhead box M1 transcription factor contributes to the development and growth of mouse colorectal cancer. Gastroenterology, 2007. 132(4): p. 1420–31.

41. Holt, S.V., et al., Silencing Cenp-F weakens centromeric cohesion, prevents chromosome alignment and activates the spindle checkpoint. J Cell Sci, 2005. 118(Pt 20): p. 4889–900.

42. Laoukili, J., et al., FoxM1 is required for execution of the mitotic programme and chromosome stability. Nat Cell Biol, 2005. 7(2): p. 126–36.

43. Liao, H., et al., CENP-F is a protein of the nuclear matrix that assembles onto kinetochores at late G2 and is rapidly degraded after mitosis. J Cell Biol, 1995. 130(3): p. 507–18.

44. Rattner, J.B., et al., CENP-F is a. ca 400 kDa kinetochore protein that exhibits a cell-cycle dependent localization. Cell Motil Cytoskeleton, 1993. 26(3): p. 214–26.

45. Missinato, M.A., et al., Dusp6 attenuates Ras/MAPK signaling to limit zebrafish heart regeneration. Development, 2018. 145(5).

46. Gonzalez-Rosa, J.M., et al., Myocardial Polyploidization Creates a Barrier to Heart Regeneration in Zebrafish. Dev Cell, 2018. 44(4): p. 433-446.e7.

47. Jopling, C., et al., Zebrafish heart regeneration occurs by cardiomyocyte dedifferentiation and proliferation. Nature, 2010. 464(7288): p. 606–9.

48. Missinato, M.A., et al., Extracellular component hyaluronic acid and its receptor Hmmr are required for epicardial EMT during heart regeneration. Cardiovasc Res, 2015. 107(4): p. 487–98.

49. Bolte, C., et al., Expression of Foxm1 transcription factor in cardiomyocytes is required for myocardial development. PLoS One, 2011. 6(7): p. e22217.

50. Camenisch, T.D., et al., Disruption of hyaluronan synthase-2 abrogates normal cardiac morphogenesis and hyaluronan-mediated transformation of epithelium to mesenchyme. J Clin Invest, 2000. 106(3): p. 349–60.

51. Ramakrishna, S., et al., Myocardium defects and ventricular hypoplasia in mice homozygous null for the Forkhead Box M1 transcription factor. Dev Dyn, 2007. 236(4): p. 1000–13.

52. Dunsch, A.K., et al., Dynein light chain 1 and a spindle-associated adaptor promote dynein asymmetry and spindle orientation. J Cell Biol, 2012. 198(6): p. 1039–54.

53. Korver, W., J. Roose, and H. Clevers, The winged-helix transcription factor Trident is expressed in cycling cells. Nucleic Acids Res, 1997. 25(9): p. 1715–9.

54. Korver, W., et al., Uncoupling of S phase and mitosis in cardiomyocytes and hepatocytes lacking the winged-helix transcription factor Trident. Curr Biol, 1998. 8(24): p. 1327–30.

55. Hatano, H., et al., Overexpression of receptor for hyaluronan-mediated motility (RHAMM) in MC3T3-E1 cells induces proliferation and differentiation through phosphorylation of ERK1/2. J Bone Miner Metab, 2012. 30(3): p. 293–303.

56. Wonsey, D.R. and M.T. Follettie, Loss of the forkhead transcription factor FoxM1 causes centrosome amplification and mitotic catastrophe. Cancer Res, 2005. 65(12): p. 5181–9.

57. Sengupta, A., V.V. Kalinichenko, and K.E. Yutzey, FoxO1 and FoxM1 transcription factors have antagonistic functions in neonatal cardiomyocyte cell-cycle withdrawal and IGF1 gene regulation. Circ Res, 2013. 112(2): p. 267–77.

58. Kettleborough, R.N., et al., A systematic genome-wide analysis of zebrafish protein-coding gene function. Nature, 2013. 496(7446): p. 494–7.

59. Liu, X., et al., Neuregulin-1/erbB-activation improves cardiac function and survival in models of ischemic, dilated, and viral cardiomyopathy. J Am Coll Cardiol, 2006. 48(7): p. 1438–47.

60. Bersell, K., et al., Neuregulin1/ErbB4 signaling induces cardiomyocyte proliferation and repair of heart injury. Cell, 2009. 138(2): p. 257–70.

61. D’Uva, G., et al., ERBB2 triggers mammalian heart regeneration by promoting cardiomyocyte dedifferentiation and proliferation. Nat Cell Biol, 2015. 17(5): p. 627–38.

62. Ma, R.Y., et al., Raf/MEK/MAPK signaling stimulates the nuclear translocation and transactivating activity of FOXM1c. J Cell Sci, 2005. 118(Pt 4): p. 795–806.

63. Lam, E.W., et al., Forkhead box proteins: tuning forks for transcriptional harmony. Nat Rev Cancer, 2013. 13(7): p. 482–95.

64. Beisaw, A., et al., AP-1 Contributes to Chromatin Accessibility to Promote Sarcomere Disassembly and Cardiomyocyte Protrusion During Zebrafish Heart Regeneration. Circ Res, 2020. 126(12): p. 1760–1778.

65. Fukuda, R., et al., Stimulation of glycolysis promotes cardiomyocyte proliferation after injury in adult zebrafish. EMBO Rep, 2020. 21(8): p. e49752.

66. Honkoop, H., et al., Single-cell analysis uncovers that metabolic reprogramming by ErbB2 signaling is essential for cardiomyocyte proliferation in the regenerating heart. Elife, 2019. 8.

67. Bomont, P., et al., Unstable microtubule capture at kinetochores depleted of the centromere-associated protein CENP-F. Embo j, 2005. 24(22): p. 3927–39.

68. Feng, J., H. Huang, and T.J. Yen, CENP-F is a novel microtubule-binding protein that is essential for kinetochore attachments and affects the duration of the mitotic checkpoint delay. Chromosoma, 2006. 115(4): p. 320–9.

69. Dees, E., et al., Cardiac-specific deletion of the microtubule-binding protein CENP-F causes dilated cardiomyopathy. Dis Model Mech, 2012. 5(4): p. 468–80.

70. Dees, E., et al., LEK1 protein expression in normal and dysregulated cardiomyocyte mitosis. Anat Rec A Discov Mol Cell Evol Biol, 2005. 286(1): p. 823–32.

71. Patterson, M., et al., Frequency of mononuclear diploid cardiomyocytes underlies natural variation in heart regeneration. Nat Genet, 2017. 49(9): p. 1346–1353.

72. Wills, A.A., et al., Regulated addition of new myocardial and epicardial cells fosters homeostatic cardiac growth and maintenance in adult zebrafish. Development, 2008. 135(1): p. 183–92.

73. Kirillova, A., et al., Polyploid cardiomyocytes: implications for heart regeneration. Development, 2021. 148(14).

74. Hesse, M., et al., Midbody Positioning and Distance Between Daughter Nuclei Enable Unequivocal Identification of Cardiomyocyte Cell Division in Mice. Circ Res, 2018. 123(9): p. 1039–1052.

75. Leone, M., G. Musa, and F.B. Engel, Cardiomyocyte binucleation is associated with aberrant mitotic microtubule distribution, mislocalization of RhoA and IQGAP3, as well as defective actomyosin ring anchorage and cleavage furrow ingression. Cardiovasc Res, 2018. 114(8): p. 1115–1131.

76. Liu, H., et al., Control of cytokinesis by β-adrenergic receptors indicates an approach for regulating cardiomyocyte endowment. Sci Transl Med, 2019. 11(513).

77. Chablais, F. and A. Jazwinska, The regenerative capacity of the zebrafish heart is dependent on TGFβ signaling. Development, 2012. 139(11): p. 1921–30.

78. Choi, W.-Y., et al., <em>In vivo</em> monitoring of cardiomyocyte proliferation to identify chemical modifiers of heart regeneration. Development, 2013. 140(3): p. 660–666.

79. Huang, Y., et al., Igf Signaling is Required for Cardiomyocyte Proliferation during Zebrafish Heart Development and Regeneration. PLoS One, 2013. 8(6): p. e67266.

80. Jabbour, A., et al., Parenteral administration of recombinant human neuregulin-1 to patients with stable chronic heart failure produces favourable acute and chronic haemodynamic responses. Eur J Heart Fail, 2011. 13(1): p. 83–92.

81. Lepilina, A., et al., A dynamic epicardial injury response supports progenitor cell activity during zebrafish heart regeneration. Cell, 2006. 127(3): p. 607–19.

82. Parodi, E.M. and B. Kuhn, Signalling between microvascular endothelium and cardiomyocytes through neuregulin. Cardiovasc Res, 2014. 102(2): p. 194–204.

83. Zhao, Y.Y., et al., Neuregulins promote survival and growth of cardiac myocytes. Persistence of ErbB2 and ErbB4 expression in neonatal and adult ventricular myocytes. J Biol Chem, 1998. 273(17): p. 10261–9.

84. Beauchemin, M., A. Smith, and V.P. Yin, Dynamic microRNA-101a and Fosab expression controls zebrafish heart regeneration. Development, 2015. 142(23): p. 4026–37.

85. van Duijvenboden, K., et al., Conserved NPPB+ Border Zone Switches From MEF2-to AP-1-Driven Gene Program. Circulation, 2019. 140(10): p. 864–879.

86. Wu, H.Y., et al., Fosl1 is vital to heart regeneration upon apex resection in adult Xenopus tropicalis. NPJ Regen Med, 2021. 6(1): p. 36.

87. Chen, X., et al., The forkhead transcription factor FOXM1 controls cell cycle-dependent gene expression through an atypical chromatin binding mechanism. Mol Cell Biol, 2013. 33(2): p. 227–36.

88. Down, C.F., et al., Binding of FoxM1 to G2/M gene promoters is dependent upon B-Myb. Biochim Biophys Acta, 2012. 1819(8): p. 855–62.

89. Sadasivam, S., S. Duan, and J.A. DeCaprio, The MuvB complex sequentially recruits B-Myb and FoxM1 to promote mitotic gene expression. Genes Dev, 2012. 26(5): p. 474–89.

90. Sanders, D.A., et al., FOXM1 binds directly to non-consensus sequences in the human genome. Genome Biol, 2015. 16(1): p. 130.

91. Wang, X., et al., The Forkhead Box m1b transcription factor is essential for hepatocyte DNA replication and mitosis during mouse liver regeneration. Proc Natl Acad Sci U S A, 2002. 99(26): p. 16881–6.

92. Yan, D., et al., Activation of AKT/AP1/FoxM1 signaling confers sorafenib resistance to liver cancer cells. Oncol Rep, 2019. 42(2): p. 785–796.

93. Warburg, O., F. Wind, and E. Negelein, THE METABOLISM OF TUMORS IN THE BODY. J Gen Physiol, 1927. 8(6): p. 519–30.

94. Makinde, A.O., P.F. Kantor, and G.D. Lopaschuk, Maturation of fatty acid and carbohydrate metabolism in the newborn heart. Mol Cell Biochem, 1998. 188(1-2): p. 49–56.

95. Nakano, H., et al., Glucose inhibits cardiac muscle maturation through nucleotide biosynthesis. Elife, 2017. 6.

96. Cui, J., et al., FOXM1 promotes the warburg effect and pancreatic cancer progression via transactivation of LDHA expression. Clin Cancer Res, 2014. 20(10): p. 2595–606.

97. Wang, Y., et al., FOXM1 promotes reprogramming of glucose metabolism in epithelial ovarian cancer cells via activation of GLUT1 and HK2 transcription. Oncotarget, 2016. 7(30): p. 47985–47997.

98. Han, L., et al., Lamin B2 Levels Regulate Polyploidization of Cardiomyocyte Nuclei and Myocardial Regeneration. Dev Cell, 2020. 53(1): p. 42-59.e11.

99. Hirose, K., et al., Evidence for hormonal control of heart regenerative capacity during endothermy acquisition. Science, 2019. 364(6436): p. 184–188.

100. Cheng, Y.Y., et al., Reprogramming-derived gene cocktail increases cardiomyocyte proliferation for heart regeneration. EMBO Mol Med, 2017. 9(2): p. 251–264.

101. Huang, C.J., et al., Germ-line transmission of a myocardium-specific GFP transgene reveals critical regulatory elements in the cardiac myosin light chain 2 promoter of zebrafish. Dev Dyn, 2003. 228(1): p. 30–40.

102. Perner, B., C. Englert, and F. Bollig, The Wilms tumor genes wt1a and wt1b control different steps during formation of the zebrafish pronephros. Dev Biol, 2007. 309(1): p. 87–96.

