## Supplementary figures and images for "Foxm1 drives cardiomyocyte proliferation in adult zebrafish after cardiac injury"

### Fig. S1

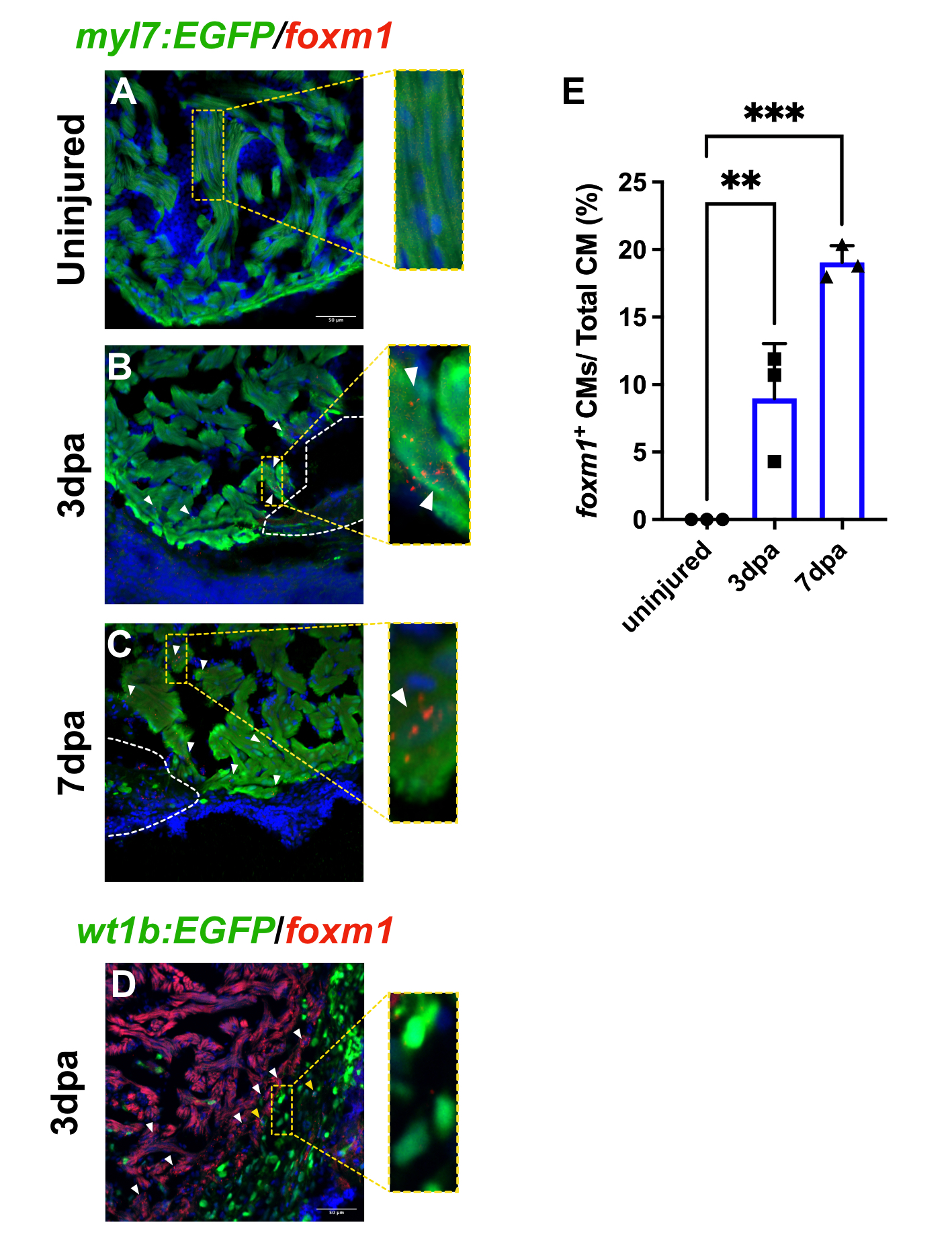

### Fig. S2

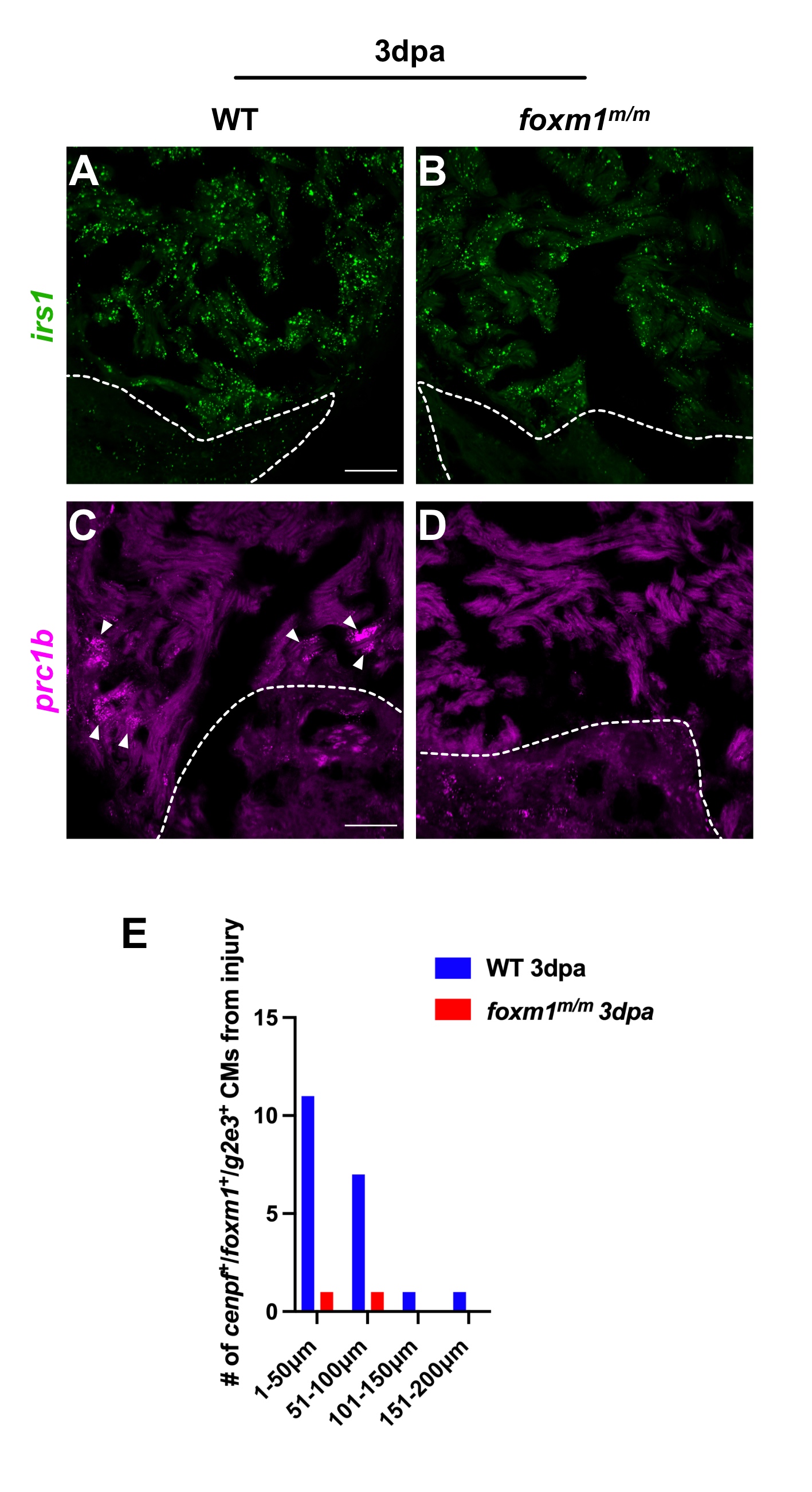

### Fig. S3

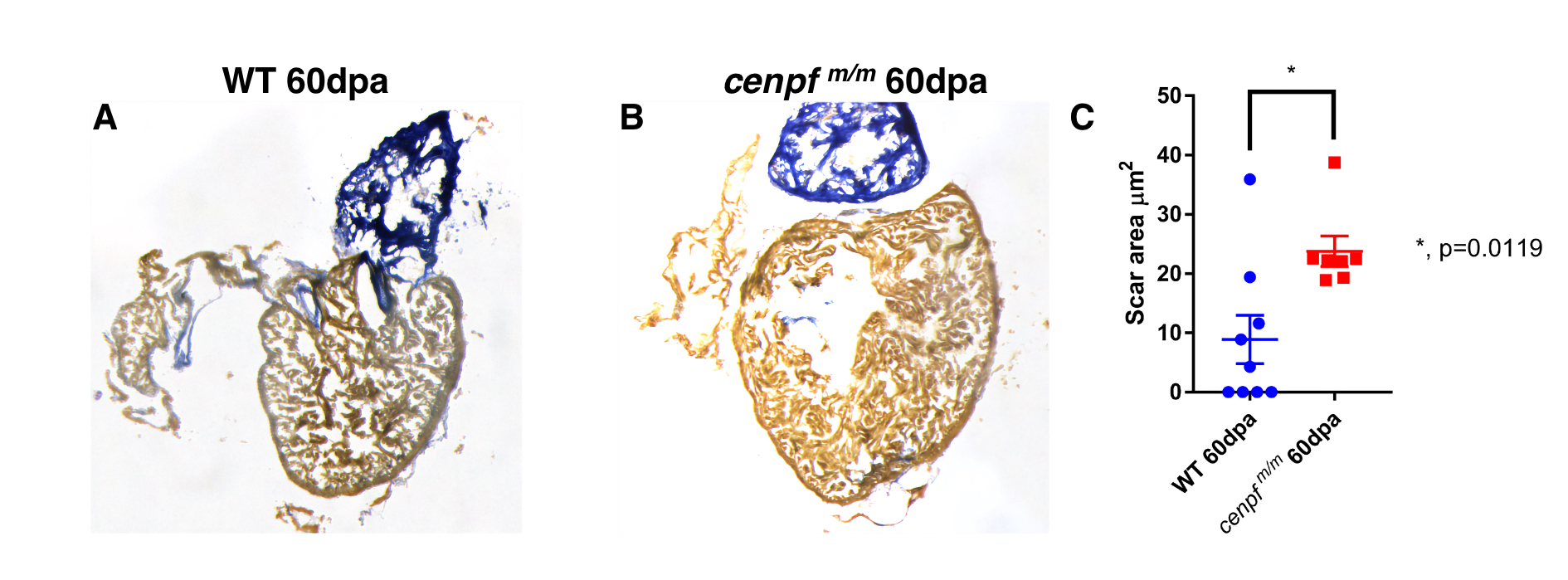
