## Supplementary Table 5 for "Foxm1 drives cardiomyocyte proliferation in adult zebrafish after cardiac injury"

**Primer sequences for Genotyping:**

| **ID** | **Gene Symbol** | **Forward Primer Sequence (5′-3′)** | **Reverse Primer Sequence (5′-3′)** |
| --- | --- | --- | --- |
| XM_002665215.6 | *cenpf* | CTAAACCAGTCTTCGGTGAGCCATAAGG | CCAACTGTGTCACTCGGTGCGCCTGATGTGAACTACT |
| NM_194380.2 | *dusp6* | GTGGCTCGCGCACTCACAGGCTA | GGAGCCGCCGTCGATGTTTTCA |
| NM_201097.1 | *foxm1* | ATCTCCTGATGGCAAGATCTCCTATTGGACGAGT | GGTCAACGGGTCACCCAAAGGCTGTGGG |
| NM_199580.2 | *hmmr* | CTTGAAATGAAACATCAGAGACTACTAGAGAAACCGG | CTTCAGTTCTTCCTCCAGGGCC |

**Table: Primer sequences for qPCR Analysis:**

| **ID** | **Gene Symbol** | **Forward Primer Sequence (5′-3′)** | **Reverse Primer Sequence (5′-3′)** |
| --- | --- | --- | --- |
| XM_001340930.6 | *alx4a* | CGCGTGTTATGGTAAAGACAGC | ACGGTTGCGTCTTTTCTTGC |
| NM_181601.5 | *bactin2* | CGTGCTGTCTTCCCATCCA | TCACCAACGTAGCTGTCTTTCTG |
| NM_001076719.1 | *ccnb3* | GGAGGATGGACGCAGTGTTTA | TGGAGTTTTCTGACCACGGG |
| NM_131025.4 | *ccnd1* | ACGCAGCTACAATCAGCAGT | GCAGCACCATGTCAAAAGGG |
| NM_207048.1 | *ccnf* | GTTTGACGATGAGGAGAAGCG | TGATTGCCCACAGCAGGTAG |
| NM_213172.1 | *ccng2* | GAAATCGCACACCGCATACA | CACTTGAATCCTCACTCTCGCTC |
| NM_213080.2 | *cdc20* | AGATATGCCCATCACCAACGCC | GGGATGTACCGATCTCCTCCTG |
| NM_212564.2 | *cdk1* | TCGCCACCAAGAAACCTCTC | AAGGTCGATGCCGTTCTTGT |
| XM_002665215.6 | *cenpf* | TTTAAAAGTTAAGACACAAACCCAGA | CCAATTCTTGAGCAGATGATACAC |
| NM_001100093.2 | *ercc6l* | CCGATAAGTTTGGGGGCTGT | TCCTTCCCCTTCTGCCGATA |
| NM_205569.1 | *fosab* | TTACCCGCTCAACCAGACTC | TGAAGAGATCGCCGTGACAG |
| NM_001007312.1 | *fosb* | GCTTTTCGTCCGGTGGTTTC | TGAGTACAGCATCGCTCACC |
| NM_201097.1 | *foxm1* | CCAAAGGTCTCTCAGAGCGACT | AGGTCACAGGCTGAAACAC |
| NM_001003822.1 | *g2e3* | GCTAAAAGGCAGTGTGTGCC | CACCAGCACCTCTGTGAACT |
| NM_001115114.1 | *gapdh* | CCGTCTTGAGAAACCTGCCAA | GGATGAACGGCAATCCCCAT |
| XM_017356649.2 | *hdac7b* | CCTTTGGAGTTTACAGCAGTGG | GTGAGACATCTGTCACGTCTG |
| NM_200405.1 | *hif1al* | GTGTGACACAGCAGCTCTCAT | GATGGCAATGTTTCCGCGAC |
| NM_213252.1 | *hk1* | AATGCAAGGAACCGGAGAGG | TTCACCAAGAGTCCCGCATC |
| NM_173283.4 | *igfbp1a* | AGCGAGACAGCACCAGATCAT | TGCGTATCGCGTTGACTTTG |
| NM_001161402.2 | *igfbp6b* | GGAAAAAGCAGTGTCGGTCC | TATCCCTGGTGTGTCCTGGT |
| XM_682610.9 | *irs1* | TAGGCTGACACGTCCAACAC | AAACAGGTGGGAACAGCGAT |
| XM_695654.9 | *irs2b* | CTTGTCCGGTTCAAAGCTGC | CAGCCGAGGACTGAAGTTGT |
| NM_001128342.1 | *jund* | CAACGACGCCATAAATCGCA | ATCCTACACTCTCCTGCCGT |
| NM_001100089.1 | *kmt5ab* | GGAAGAGGGGTTTTTGCGGA | TGGTAGCGTCCACACAGTAG |
| NM_131616.3 | *pbx3b* | GTATCCAACTGGTTCGGCAA | AGGGTATCCACCGGAGTTGG |
| NM_198816.1 | *pfkfb4b* | CTGCGGAAGATATGGATGCC | GCCGACCGTCACAATGAGAG |
| XM_005162525.4 | *pfkpa* | GACCGTAACTTCGGGACCAA | GCGAACACTCTCCCTTCATCA |
| NM_001003488.1 | *pkmb* | GGGCTTATTAAGGGCAGTGG | CCAACATGCCTCCGTTCTCA |
| NM_001002317.2 | *polr2d* | CCAGATTCAGCCGCTTCAAG | CAAACTGGGAATGAGGGCTT |
| NM_131112.1 | *pou5f3* | AGAGAGATGTAGTGCGTGTATGG | CATGTATAAGGCAGGGGCTC |
| NM_200234.1 | *prc1b* | ATGGCACTGTAATCCGCACA | ACCCCATTACCCTTGCTTGC |
| NM_001245966.1 | *zeb2b* | ACACGAGGAAAACGACCTGC | GCTGAGTGCGGTAAGCAAAC |
